# Looking at the BiG picture: Incorporating bipartite graphs in drug response prediction

**DOI:** 10.1101/2021.08.11.455993

**Authors:** David Earl Hostallero, Yihui Li, Amin Emad

## Abstract

**Motivation:** The increasing number of publicly available databases containing drugs’ chemical structures, their response in cell lines, and molecular profiles of the cell lines has garnered attention to the problem of drug response prediction. However, many existing methods do not fully leverage the information that is shared among cell lines and drugs with similar structure. As such, drug similarities in terms of cell line responses and chemical structures could prove to be useful in forming drug representations to improve drug response prediction accuracy.

**Results:** We present two deep learning approaches, BiG-DRP and BiG-DRP+, for drug response prediction. Our models take advantage of the drugs’ chemical structure and the underlying relationships of drugs and cell lines through a bipartite graph and a heterogenous graph convolutional network that incorporate sensitive and resistant cell line information in forming drug representations. Evaluation of our methods and other state-of-the-art models in different scenarios shows that incorporating this bipartite graph significantly improves the prediction performance. Additionally, genes that contribute significantly to the performance of our models also point to important biological processes and signaling pathways. Analysis of predicted drug response of patients’ tumors using our model revealed important associations between mutations and drug sensitivity, illustrating the utility of our model in pharmacogenomics studies.

**Availability and Implementation:** An implementation of the algorithms in Python is provided in github.com/ddhostallero/BiG-DRP.

**Supplementary Information:** Online-only supplementary data is available at the journal’s website.

## INTRODUCTION

Utilization of machine learning and statistical analyses in precision medicine has gained attention in the past decade. Prediction of drug response based on samples’ molecular profiles is a major problem in this domain and various approaches have been proposed for this purpose [1–5]. Gene expression profile of samples is widely used for this purpose due to their higher predictive ability compared to other molecular profiles [1]. The curation of large public databases of gene expression profiling of hundreds of cancer cell lines (CCLs) and their response to hundreds of different drugs (e.g., GDSC [6]) has accelerated the development of novel methodologies in this domain.

Due to the similarity in molecular and chemical structure of different drugs and their mechanisms of action, machine learning (ML) methods that can take advantage of these similarities are of great interest. Instead of training a different ML model for each drug, one can formulate the drug response prediction as a paired prediction problem, such that a model takes in a (drug, CCL) pair as input and trains a single model for all drugs and CCLs [7–9]. This increases the number of samples, and enables information sharing across many drugs and drug families. Chemical structure data (e.g., PubChem [10], ChEMBL [11]) is particularly useful for representing the drugs, and models have been developed to take advantage of these [12–14].

Some approaches [15–17] have formulated this as a matrix factorization problem, forming a matrix of drugs and CCLs. One advantage of this is that these methods directly work with the “entities” (i.e., drugs and CCLs) and responses, and do not need to map feature representations of the entities to their responses, although available features can be utilized for regularization [15, 17]. However, this formulation is inherently transductive, since samples and drugs are expected to be present in the matrix. As a result, these models cannot be directly used to predict the response of a new CCL to a drug unless the CCL has drug response information in the training set for some other drugs prior to training. Another group of methods utilize collaborative filtering [18, 19] and predictions are calculated using an entity’s neighborhood, which are defined by the similarities calculated from gene expressions, molecular fingerprints, and drug responses. Since these approaches require the calculation of drug response similarities, an inherent assumption is to have at least a few known responses for each unique CCL and drug in the test set, which is a more relaxed assumption compared to that of matrix factorization methods.

Taking inspiration from the concept of “entity” from the matrix factorization approaches and to overcome their shortcoming due to their transductive nature, we propose to utilize the underlying matrix by transforming these entities into drug and CCL nodes and form a bipartite graph. We hypothesized that incorporating cell line information that are highly sensitive or resistant to a drug could improve the drug representation for drug response prediction. In our approach called Bipartite Graph-represented Drug Response Predictor (BiG-DRP and BiG-DRP+), we formed this graph by filtering the most sensitive and resistant CCLs for each drug, and linking them through an edge. Although drugs are not directly connected to each other through an edge, 2-hop message passing incorporates information on drug similarities. The model accepts drugs’ descriptors and CCLs’ gene expression profiles as input, and utilizes them as node attributes for the bipartite graph and as sample features. The output is a continuous drug response value pertaining to the predicted normalized log IC50.

To evaluate the performance of BiG-DRP and BiG-DRP+, we used 5-fold cross validation and compared these results across different baselines and other drug response prediction approaches, namely NRL2DRP [20], PathDNN [7], and tCNN [9]. We tested on two data-splitting methods, 5-fold leave-pairs-out and 5-fold leave-cell lines-out, which represent two possible scenarios of data availability. In both scenarios, we have shown significant improvement compared to other approaches. In addition, using a computational pipeline that we developed for identifying the most contributing features, we identified genes that pointed to biological processes and signaling pathways involved in drugs’ mechanisms of action.

## METHODS

### Bipartite Graph-based Drug Response Prediction

We developed a novel deep learning-based drug response prediction model that takes advantage of a bipartite graph between drugs and cell lines, which we called Bipartite Graph-represented Drug Response Predictor (BiG-DRP). We also proposed an extension of BiG-DRP, called BiG-DRP+, which accounts for constantly changing drug representations in the former approach. An overview of the (shared) architecture of these models are provided in Figure 1.

**Figure 1:**
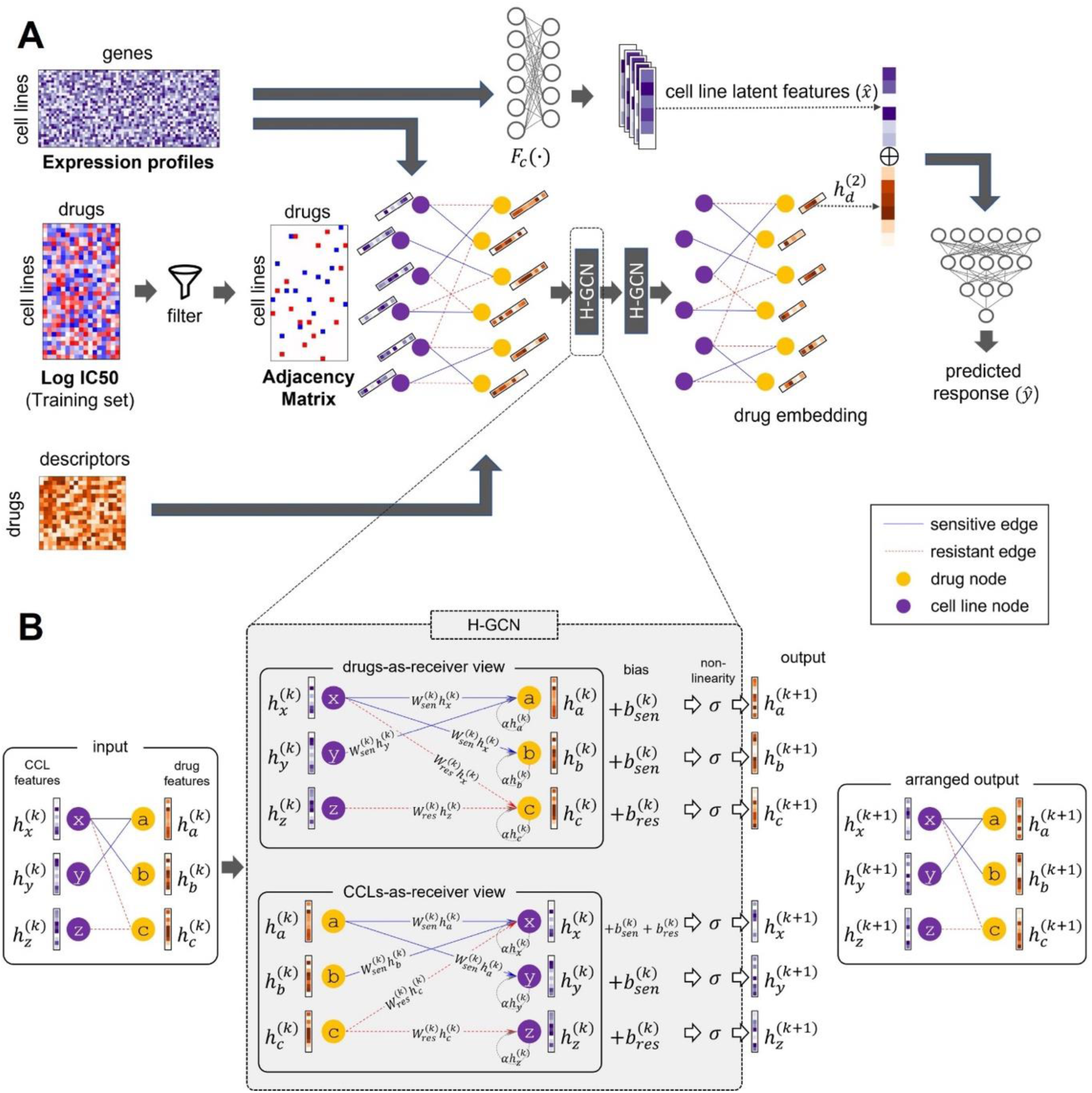
The computational pipeline and architecture of BiG-DRP and BiG-DRP+. A) Latent drug embeddings are generated using a heterogenous graph convolutional network based on a bipartite graph of drug-CCLs and drug descriptors. In parallel, CCL embeddings are generated using an encoder neural network based on their gene expression profile. These embeddings are then used by a predictor neural network to predict the drug response values. B) An overview of a single H-GCN layer is shown (our models use two stacked H-GCN layers). The H-GCN propagates information to neighbouring nodes by taking the graph as an input. Node attributes are multiplied to the weight matrices (***W_sen_*** and ***W_res_***) to produce the messages, which will be sent to their neighbours, depending on the type of edge between two nodes. Each of the nodes will then aggregate their received messages, along with the biases and self-information. The output is then the same graph, whose nodes’ attributes include information from neighbouring nodes and their connectivity.

The BiG-DRP pipeline first obtains latent embeddings of CCLs and drugs and uses them in the drug response prediction task. To obtain drug embeddings, first a heterogeneous bipartite graph composed of CCL nodes and drug nodes is formed. The nodes of the bipartite graph are connected via two types of edges: sensitive edges or resistant edges. These edges are based on the log IC50 values of each CCL-drug pair. A sensitive edge implies that the CCL is likely to be sensitive to the drug, while a resistant edge implies that it is likely to be resistant to the drug. In addition, each CCL node is assigned attributes corresponding to its gene expression (GEx) profile and each drug node is assigned attributes corresponding to its drug descriptors. Then, a heterogenous graph convolutional network (H-GCN) generates embeddings of each drug (denoted as *h_d_*^(2)^ in Figure 1) using this bipartite graph. For each drug of interest, the H-GCN obtains an embedding that not only captures the molecular characteristic of the drug itself, but also captures the characteristics of other drugs that induce a similar sensitive/resistant pattern in CCLs. Inclusion of the GEx profiles of CCLs as node attributes in the bipartite graph allows the model to define the “similar pattern” mentioned above in a broader sense: instead of requiring a similar pattern in the exact same CCLs, the model can identify such patterns in CCLs that have a similar GEx profile.

To obtain embeddings of the CCLs based on their GEx profiles (denoted as 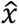 in Figure 1), the model uses a neural network that is separate from the H-GCN. While it is possible to use the bipartite graph and the H-GCN to obtain CCL embeddings, such a choice would limit the applicability of the pipeline to only CCLs that are already present in the training set. The reason is that a CCL that is not present in the training set will be in the form of a single disconnected node in the bipartite graph and no embedding can be found for it using the H-GCN. However, in many practical applications (e.g., prediction of clinical drug response of patients based on models trained on preclinical CCLs [4, 5]), a model must be able to predict drug response of samples that are not seen by the model during the training for any drug. To avoid this limitation, the CCL embeddings are obtained independent of the H-GCN network and the bipartite graph. The drug and CCL embeddings are then concatenated, representing each (drug, CCL) pair. Then, a series of neural network layers (collectively called the *predictor*) are used to predict the drug response of each such pair using the concatenated embeddings.

The BiG-DRP+ is an extension of BiG-DRP with the exact same architecture, which aims to stabilize the trained model. After the “last” training epoch of BiG-DRP (i.e., starting from BiG-DRP’s trained weights), we train the model for one more epoch but with a smaller learning rate and “frozen” drug embeddings. The lower learning rate prevents the predictor from overfitting while the freezing of the embeddings allows the predictor to learn the finite set of drugs instead of constantly changing representations of the exact same drugs.

### Construction of the Heterogenous Bipartite Graph

We denote the heterogenous bipartite graph as *G*(*V_C_*, *V_D_*, *E_r_*, *E_s_*), where *V_C_* is the set of CCL nodes used to build the graph (a subset of all the CCLs in the study) and *V_D_* is the set of drug nodes. *E_r_* is the set of edges that connect drugs to their “most resistant” CCLs, while *E_s_* is the set of edges that connect drugs to their “most sensitive” CCLs. For a fixed value of *k*, a drug is connected via a resistant edge to CCLs whose log IC50 is among the top *k* percent and is connected via a sensitive edge to CCLs whose log IC50 is among the bottom *k* percent of the CCLs. The set *V_C_* is then the union of all such CCLs whose drug response are among the top *k* or bottom *k* percent of all cell lines for at least one drug. It is worth noting that the edges in this graph are unweighted and the log IC50 values are only used to determine whether a resistant (or sensitive) edge exists or not. We used *k* = 1 in our analysis, but the performance of BiG-DRP and BiG-DRP+ were not sensitive to the choice of *k*, as discussed in Results.

### Drug embedding using heterogenous graph convolutions

We used a 2-layer heterogenous graph convolutional network (H-GCN) to find a network-based embedding of the drugs. An H-GCN is a variation of graph convolutional network [21], which allows multiple edge types. A forward pass of an H-GCN can be summarized using the following equation:

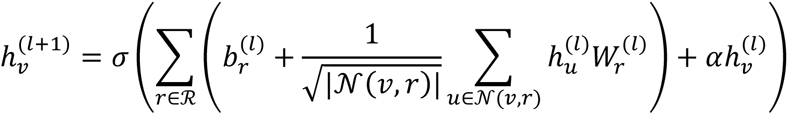

where 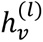 is node 𝑣’s embedding at the *l*th layer, *σ* is a non-linearity function, 𝒩(𝑣, *r*) is node 𝑣’s set of neighbours connected using the edge type *r*. 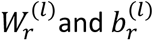 are the weights and biases at the *l*th H-GCN layer for edge type *r*, respectively. Intuitively, this allows a separation of GCN parameters for each edge type, and thus creates context during the message passing. The normalization factor 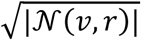 prevents the embedding values from exploding due to a large number of neighbours.

Although we constructed a bipartite graph, artificially adding self-loops to the graph is a common practice in GCNs to retain some information from the previous layer, avoiding the complete dependence of the node’s embedding to its neighbours. However, in the case of H-GCN, self-loops increase the complexity of the model by adding another set of parameters. To avoid this, we injected a residual term 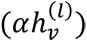 to the forward pass to simulate self-loops. Here, *α* is a hyperparameter (we fixed the value to *α* = 0.5 ) pertaining to the amount of information to be retained for the next layer.

The bipartite graph and the H-GCN allow us to find a drug embedding that captures relevant information about the CCLs that are generally resistant/sensitive to it (its 1-hop neighbours), as well as information on other drugs to which these CCLs have a similar or inverse pattern of response (its 2-hop neighbours). These embeddings enable sharing of information across drugs that are connected to similar set of cell lines via similar edge types.

### Data Acquisition and Preprocessing

We obtained the drug response data in the form of log IC50 values from the Genomics of Drug Sensitivity in Cancer (GDSC) database [6]. We only selected drugs with known log IC50 values as well as binarized responses that allow us to calculate the key performance metrics used for evaluation of different methods. We also filtered out duplicate drugs that came from different batches, which are tagged with different drug IDs, named with synonyms, or labeled as “rescreens”. In cases of such duplicates, we only kept the one for which the drug response in a larger number of cell lines was measured. We collected the Simplified Molecular Input Line Entry System (SMILES) encoding [22] of these drugs and used the RDKit software [12] to generate drug descriptors (e.g. molecular weight, number of aromatic rings) from these encodings. Descriptors with missing values were excluded from the analysis. At the end of these data cleaning steps, we were left with 237 unique drugs, each with feature vectors of length 198 (representing their drug descriptors).

We performed z-score normalization on drug descriptors, one feature at a time across all drugs. We also z-score normalized the log IC50 values of each drug (one drug at a time) across cell lines. This is necessary since the log IC50 values of different drugs have significantly different means and standard deviations, which renders the calculated metrics incomparable across drugs and inflates the overall correlation coefficient. For example, a relatively small mean squared error for a certain drug, or a high overall spearman correlation do not necessarily indicate good performance without such a normalization. This drug-wise normalization allows us to compare results across different drugs, and prevents overestimation of the models’ performance.

For the 237 drugs above, we obtained the RNA-seq GEx profile of 1001 CCLs from the Cell Model passports [23]. We performed log_2_(FPKM +1) transformation on the FPKM values. We excluded genes that showed a small variability across the cell lines (genes with standard deviation <0.1) as well as genes with missing values in some cell lines. After these preprocessing steps, we ended up with 944 unique CCLs and their GEx values of 13,823 genes. This amounted to a total of 181,380 labeled CCL-drug pairs.

### Training Procedure

As discussed earlier, to enable the model to generalize to completely new CCLs (those that are not seen by the model for any drug during training), we used a separate neural network, parallel to the H-GCN. As input, this network accepts the CCLs’ gene expression vector *x* and produces a latent representation 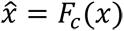. We then concatenate 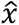 with the drug *d*’s embedding, 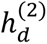, and use it as input for our predictor, a 3-layer neural network that outputs the predicted drug response values 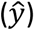.

The model was trained end-to-end using the mean squared error 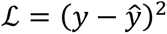 and Adam as the optimizer [24]. We also fixed the learning rate to 0.0001 and batch size to 128 (see Results for the effect of different choices of hyperparameters on the performance). We used Leaky ReLU for all non-linearity functions (i.e. *σ*(*x*) = max(0, *x*) + 0.01 × min (0, *x*)). The number of training steps were decided by randomly selecting samples from the training data and using them as a validation set for early stopping. The model was then re-trained with the entire training set and the previously identified optimal number of training steps. For BiG-DRP+, the extra epoch’s learning rate was set to 0.00001.

In our approach, elements of a batch are (drug, CCL) pairs, although all drug embeddings can be generated simultaneously for each forward pass. Embeddings generated using graph convolutional networks rely on the node connectivity. This generally means that a small perturbation of a node’s embedding may affect the embeddings of its neighbours in the next GCN (or H-GCN) layer. Unlike regular dense neural networks, it is possible that a dramatic change would occur in the embeddings, even with a relatively small learning rate. In such cases, the predictor may not easily map the “new” embedding to the “known” ones, especially if the drug was not part of the batch during the previous training step. The predictor could see this as having an infinite number of drugs, increasing the level of complexity to the learning process. To address this “moving embedding problem”, we developed BiG-DRP+, which slightly modifies the training of BiG-DRP.

The idea of BiG-DRP+ is to stop the training of the H-GCN component after several epochs but continue the training of the predictor using the “frozen” drug embeddings. In our BiG-DRP+ model, we froze the drug embeddings obtained by BiG-DRP (after the number of epochs determined by early stopping), but continued the training of other components of the architecture for one extra epoch (we used a lower learning rate for this epoch). This stabilizes the training of the predictor and enables it to identify CCLs that were treated by the same drug (since the half of the input to the predictor pertaining to the drug features are now fixed). The lower learning rate is a preventative measure to avoid overfitting.

### Evaluation and Cross-validation

To evaluate the performance of our model we used 5-fold cross validation (CV), in which one fold was kept aside as the test set for evaluation and was not used during training of the model nor for the selection of hyperparameters. This process was repeated five times (each time with a different fold as the test set) to ensure that the specific choice of the test set does not bias the results. We adopted two types of data splitting techniques to form the folds, namely leave-pairs-out (LPO) and leave-cell lines-out (LCO).

In the LPO-CV, the disjoint folds were randomly selected from the set of all (CCL, drug) pairs, while in the LCO-CV the folds contained randomly selected sets of mutually exclusive CCLs. Prior to training, GEx values were z-score normalized per gene. We used only the training folds’ unique CCLs to calculate the means and standard deviations to prevent data leakage between training and test sets. Imposing the uniqueness criterion above ensures that the models are not biased towards CCLs that exists in a larger number of (drug, CCL) pairs. To ensure a fair comparison, identical folds were used for all methods. For each drug, predictions of the five folds on their respective test sets were collected and were used to evaluate different methods.

To assess generalizability of our models to independent datasets, we obtained GEx profile (in the form of FPKM) of cancer tumours and their RECIST clinical drug response from The Cancer Genome Atlas (TCGA) [25]. Similar to previous studies [26], we considered “stable disease” or “progressive disease” as resistant and “complete response” or “partial response” as sensitive. We selected cisplatin (n = 398), paclitaxel (n = 233), gemcitabine (n = 226), and doxorubicin (n = 208), since they were present in our training dataset, had a large number of samples with known clinical drug response, and had more than 50 samples in each category of resistant and sensitive. Similar to the preprocessing steps used for GDSC dataset, the expression of each gene in the testing set (in the form of log_2_(FPKM +1)) was z-score normalized across the samples. We used PyCombat [27] to reduce the statistical discrepancies between the GDSC and TCGA samples.

### Baseline Methods

We compared our method against several baseline algorithms including both deep learning-based and traditional machine learning methods, detailed below. First, we used a multilayer perceptron (MLP) with a similar architecture and hyperparameters as BiG-DRP. Similar to BiG-DRP, the MLP also utilized the GEx and drug features. However, instead of an H-GCN, we replaced it with a dense neural network, which takes the drug features as input. We also used a support vector regressor (SVR) with a linear kernel as well as a SVR with a radial basis function (RBF) as traditional ML baselines. The concatenation of the GEx and drug features were used as the input to SVR models. Due to the large size of the data, we used Nystroem’s transformation [28] to approximate the SVR’s kernels to improve its efficiency. Hyperparameters, namely the number of Nystroem components, regularization factor, and gamma for RBF were selected using nested cross validation. In addition to the SVR models above, we used recursive feature elimination (RFE) [29] to identify the most relevant features to be used with the linear and non-linear SVR models.

NRL2DRP [20] is a graph representation learning-based method that uses a graph composed of genes, drugs, and CCL nodes, connected by edges according to their sensitivity, mutation, and protein-protein interactions. However, NRL2DRP uses a topology-based graph embedding called LINE [30], which is typically used for transductive learning. We slightly modified NRL2DRP to predict continuous values instead of binary values (so that it can be applied to our data). PathDNN [7] is another deep learning method that proposes to add some level of explainability to the drug response prediction problem by constraining the neural network connectivity using a pathway mask. This method uses drug targets and gene expressions, both of which should be in any of the Kyoto Encyclopedia of Genes and Genomes (KEGG) pathways [31]. We obtained the drug targets and pathway information from the PathDNN’s repository. The drug targets were represented by their normalized STITCH [32] confidence score, which indicates a non-zero value for genes in the drug’s targets. However, we removed three compounds because they did not have known targets in the KEGG pathways. Another deep learning approach is tCNN [9], which utilizes 1-dimensional CNNs. The canonical SMILES string of the compound is encoded into a sequence of one-hot vectors, each of which represents a single character. Since the SMILES strings vary in length, the resulting binary encoding is padded by zeros to the right to match the length of the longest encoding, resulting in a matrix of size *m* × *n*, where *m* is the number of unique characters and *n* is the length of the longest encoding. Mutations and copy number alterations, which GDSC dubs as “genetic features,” were used as the features of the CCLs.

In order to ensure a fair comparison, in our cross-validations we fixed the folds and used identical folds for each method. In addition, when an algorithm required extra information that was not used in Big-DRP, we provided those datasets as inputs to the baseline models, following the descriptions provided in each method’s manuscript. This was done to ensure we give each baseline model a fair chance.

### Identification of genes that are most predictive of drug response

To identify genes that are predictive of drug response, we used a neural network explainer called CXPlain [33] and a similar approach which we previously developed to aggregate contribution across CCLs and identify top contributors [5]. CXplain uses Granger’s causality [34] as the basis of the feature attribution. Intuitively, for each of the features, it tries to predict the increase in the sample’s loss if that specific feature is zeroed-out. We trained separate explainers for each of the drugs, since this eliminates the unnecessary complexity of learning attributions for multiple drugs, as well as the additional feature dimensions (i.e., drug features). We pooled the scores by calculating the mean attribution across all the CCLs for each of these drugs. The top genes were identified by a threshold calculated using kneedle [35], with sensitivity S=2.

### Pathway characterization analysis

We used KnowEnG’s gene set characterization pipeline [36] to perform pathway enrichment analysis (using Reactome pathways [37]). The p-values of Fisher’s exact test were corrected for multiple tests (i.e., multiple pathways) using Benjamini-Hochberg false discovery rate (FDR).

### Analysis of TCGA tumor mutations and their relationship with predicted drug responses

From TCGA database, we selected primary tumor samples that had both GEx profiles and mutation data (n = 9067). We utilized BiG-DRP+ to predict response of 237 drugs for each of the tumor samples using their GEx as input (see the Evaluation and Cross-validation section). Using the Mutation Annotation Format (MAF) file, a binary matrix indicating the existence of a mutation for a sample was formed. Similar to previous studies [38], we focused on four types of mutations: nonsense, missense, frameshift insertions and frameshift deletions. Only mutations that exist in at least 10% of the samples were included in the analysis.

## RESULTS

### Performance of BiG-DRP and BiG-DRP+ based on leave-pair-out cross validation

First, we evaluated BiG-DRP, BiG-DRP+, and other baseline algorithms using a five-fold LPO-CV, in which the folds were randomly selected among the set of all possible (CCL, drug) pairs. Table 1 shows a summary of the performance results using area under the receiver operating characteristic curve (AUROC), root mean squared error (RMSE), Pearson’s correlation coefficient (PCC) and Spearman’s correlation coefficient (SCC). To calculate these metrics across all drugs, we first calculated them separately for each drug (Supplementary Table S1) and then obtained mean and standard deviation across the drugs. BiG-DRP+ outperforms all other methods according to all metrics, and BiG-DRP outperforms all baselines but has a slightly worse performance compared to BiG-DRP+. BiG-DRP+ has a ∼5% higher AUROC and ∼11% higher SCC and PCC compared to that of MLP, which utilizes a similar architecture to BiG-DRP+ (except for the usage of the bipartite graph and the H-GCN). This highlights the importance of this novel aspect of the algorithm.

**Table 1:** The performance of BiG-DRP, BiG-DRP+ and baseline methods using 5-fold LPO-CV evaluation. Best performance values are in bold-face and underlined. The mean and standard deviations are calculated across the drugs. *Since PathDNN requires availability of at least one drug target in any of the signaling pathways, we could only apply it to 234 drugs.

| | Drug Attributes | Other inputs | Num. Drugs | AUROC mean (± std.) | RMSE mean (± std.) | SCC mean (± std.) | PCC mean (± std.) |
| --- | --- | --- | --- | --- | --- | --- | --- |
| **BiG-DRP+** | Descriptors | GEx | 237 | <u>**0.878**</u> ($\pm 0.068$) | <u>**0.843**</u> ($\pm 0.241$) | <u>**0.748**</u> ($\pm 0.100$) | <u>**0.758**</u> ($\pm 0.102$) |
| **BiG-DRP** | Descriptors | GEx | 237 | 0.875 ($\pm 0.068$) | 0.855 ($\pm 0.244$) | 0.742 ($\pm 0.099$) | 0.752 ($\pm 0.102$) |
| **Inverted BiG-DRP** | Descriptors | GEx | 237 | 0.862 ($\pm 0.075$) | 0.888 ($\pm 0.253$) | 0.721 ($\pm 0.110$) | 0.730 ($\pm 0.110$) |
| **MLP** | Descriptors | GEx | 237 | 0.835 ($\pm 0.083$) | 0.954 ($\pm 0.273$) | 0.675 ($\pm 0.120$) | 0.681 ($\pm 0.119$) |
| **NRL2DRP** | None | Drug-CCL-Gene network | 237 | 0.804 ($\pm 0.085$) | 1.153 ($\pm 0.345$) | 0.516 ($\pm 0.119$) | 0.514 ($\pm 0.123$) |
| **tCNN** | SMILES<br>One-hot<br>encoding | Genetic<br>Features | 237 | 0.787<br>($\pm 0.082$) | 1.086<br>($\pm 0.336$) | 0.587<br>($\pm 0.119$) | 0.591<br>($\pm 0.117$) |
| **PathDNN** | Drug<br>Targets | GEx,<br>pathway<br>information | 234* | 0.766<br>($\pm 0.083$) | 1.165<br>($\pm 0.355$) | 0.516<br>($\pm 0.115$) | 0.529<br>($\pm 0.117$) |
| **SVR-RBF**<br>**(w/RFE)** | Descriptors | GEx | 237 | 0.738<br>($\pm 0.101$) | 1.182<br>($\pm 0.384$) | 0.503<br>($\pm 0.125$) | 0.500<br>($\pm 0.130$) |
| **SVR-Linear**<br>**(w/RFE)** | Descriptors | GEx | 237 | 0.738<br>($\pm 0.101$) | 1.181<br>($\pm 0.393$) | 0.498<br>($\pm 0.130$) | 0.497<br>($\pm 0.135$) |
| **SVR-RBF** | Descriptors | GEx | 237 | 0.737<br>($\pm 0.100$) | 1.182<br>($\pm 0.383$) | 0.502<br>($\pm 0.123$) | 0.499<br>($\pm 0.129$) |
| **SVR-Linear** | Descriptors | GEx | 237 | 0.736<br>($\pm 0.101$) | 1.184<br>($\pm 0.393$) | 0.494<br>($\pm 0.129$) | 0.493<br>($\pm 0.134$) |

As mentioned earlier, in our models the H-GCN is used to obtain drug representations and a separate encoder is used to obtain cell line representations. We were interested to determine how the performance of the models change if we substitute the role of these two components: use the encoder to obtain drug embeddings and use the H-GCN to obtain cell line embeddings (called inverted BiG-DRP, henceforth). Our analysis showed that inverted Big-DRP outperforms all baselines, except for BiG-DRP and BiG-DRP+ (Table 1). However, it is important to note that inverted BiG-DRP has two shortcomings compared to BiG-DRP and BiG-DRP+. First, it cannot be used to predict the response of a new CCL (i.e., it cannot be used in the LCO framework), since a new CCL would not be part of the bipartite graph and as a result a representation for it cannot be obtained. Second, the bipartite graph used in inverted BiG-DRP connects each CCL to most sensitive and most resistant drugs and as a result is less reliable than the bipartite graph of BiG-DRP (that connects each drug to CCLs that are most sensitive or resistant to it). The reason is that log IC50 of different drugs for the same CCL are not directly comparable and making a bipartite graph based on this criterion may introduce errors in the network.

Figure 2 compares the performance of BiG-DRP+ against other methods for individual drugs (measured based on SCC). Each circle in the scatter plots reflects a drug, and the color of the circles reflect the density of other circles in their vicinity. Comparing BiG-DRP+ and BiG-DRP shows that the drug-specific SCC values are generally close to each other (concentrated around the diagonal line); however, the one-sided Wilcoxon signed rank test (p=2.26E-36) suggests that the performance for the majority of the drugs have improved in BiG-DRP+, albeit a small amount. Comparing to other baselines, the figure shows that the majority (and in many cases all) of the circles are above the diagonal line, suggesting a substantial improvement of their response prediction by BiG-DRP+. One-sided Wilcoxon signed rank tests also confirmed this observation, resulting in statistically significant p-values (Figure 2 and Supplementary Table S2).

**Figure 2:**
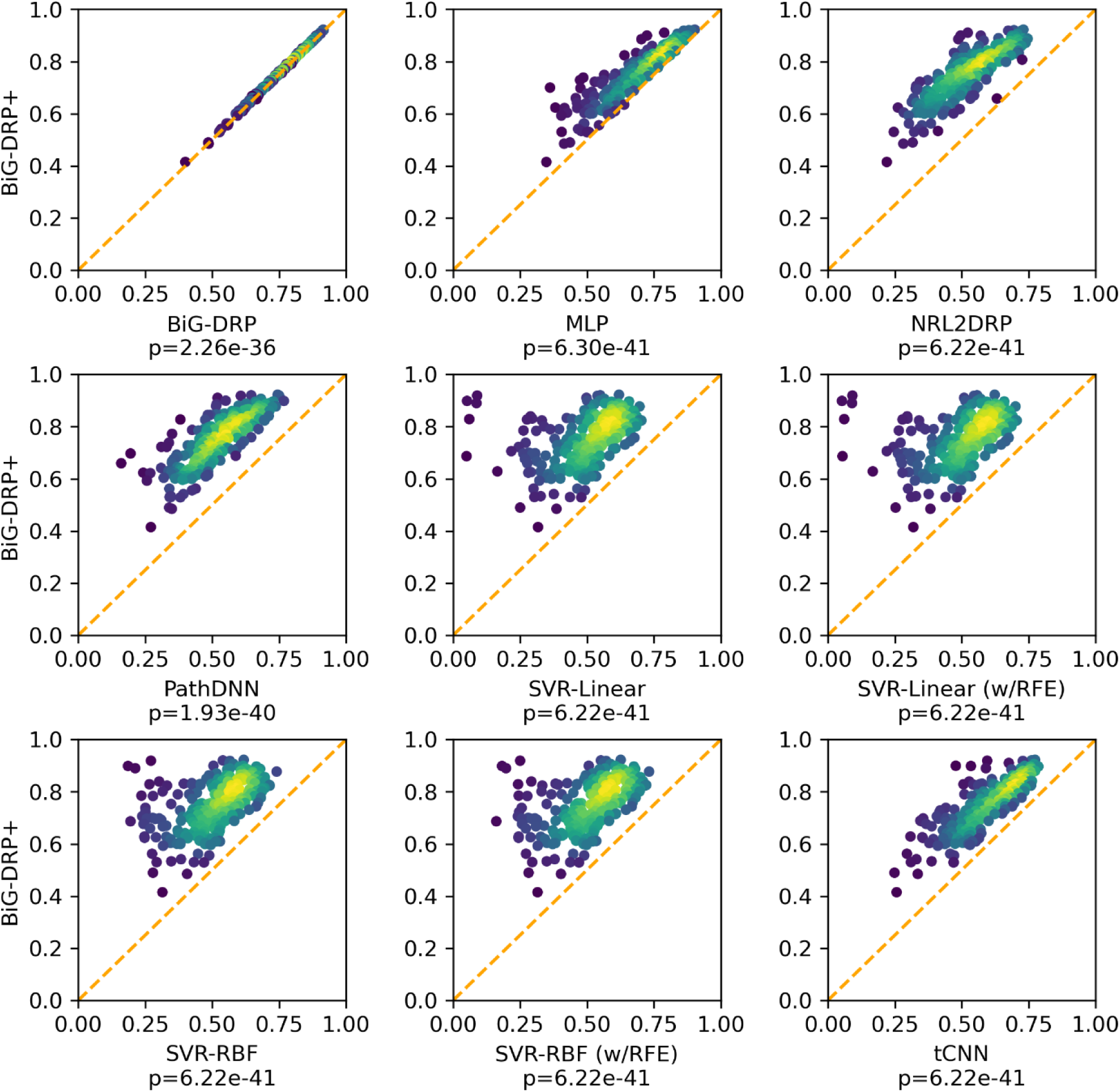
The drug-wise performance of BiG-DRP+ compared to baseline methods using five-fold LPO-CV evaluation. Each circle represents a drug, and the color of the circles reflect the density of other circles in their vicinity (yellow shows that there are many circles concentrated in that area). The coordinates reflect SCC for BiG-DRP+ (y-axis) and baseline methods (x-axis). The p-values are obtained using a one-sided Wilcoxon signed rank test, comparing the SCC of BiG-DRP+ and the baselines across drugs.

### Performance of BiG-DRP and BiG-DRP+ based on leave-cell line-out cross validation

Next, we evaluated the performance of different models using a five-fold LCO-CV. This is a stricter evaluation, since unlike LPO-CV, a CCL in the test set is never seen by the models during training, since folds are randomly selected based on the CCLs and not based on (CCL, drug) pairs. Table 2 shows the summary of the results using our performance metrics. Note that due to the transductive nature of NRL2DRP’s embedding method (LINE [30]), this method could not be applied to the LCO-CV evaluation and hence is not included in this table.

**Table 2:**
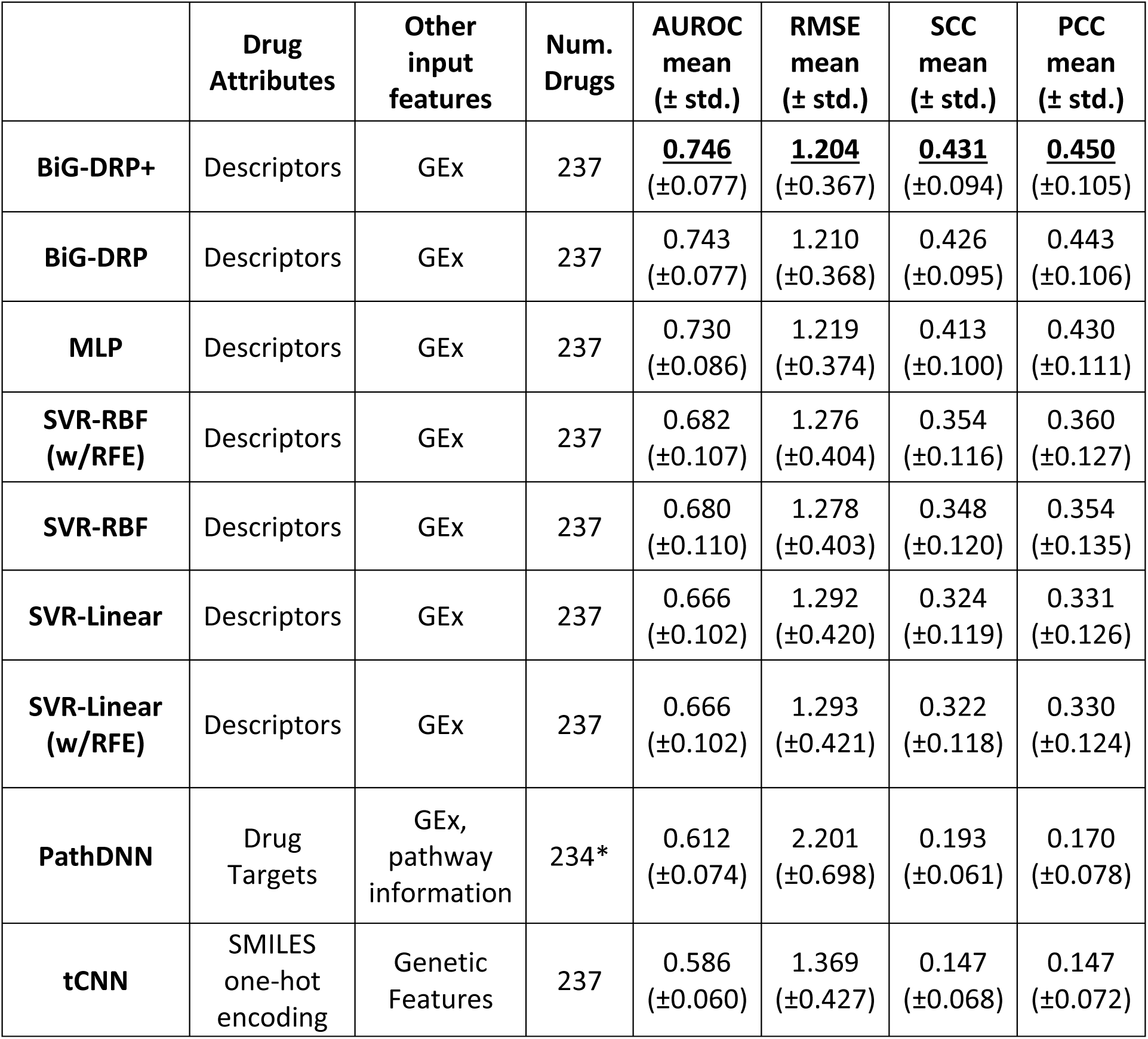
The performance of BiG-DRP+, BiG-DRP and baseline methods using five-fold LCO-CV evaluation. Best performance values are underlined. The mean and standard deviations are calculated across the drugs. *Since PathDNN requires availability of at least one drug target in any of the signaling pathways, we could only apply it to 234 drugs.

| | Drug Attributes | Other input features | Num. Drugs | AUROC mean (± std.) | RMSE mean (± std.) | SCC mean (± std.) | PCC mean (± std.) |
| --- | --- | --- | --- | --- | --- | --- | --- |
| **BiG-DRP+** | Descriptors | GEx | 237 | <u>0.746</u> ($\pm 0.077$) | <u>1.204</u> ($\pm 0.367$) | <u>0.431</u> ($\pm 0.094$) | <u>0.450</u> ($\pm 0.105$) |
| **BiG-DRP** | Descriptors | GEx | 237 | 0.743 ($\pm 0.077$) | 1.210 ($\pm 0.368$) | 0.426 ($\pm 0.095$) | 0.443 ($\pm 0.106$) |
| **MLP** | Descriptors | GEx | 237 | 0.730 ($\pm 0.086$) | 1.219 ($\pm 0.374$) | 0.413 ($\pm 0.100$) | 0.430 ($\pm 0.111$) |
| **SVR-RBF (w/RFE)** | Descriptors | GEx | 237 | 0.682 ($\pm 0.107$) | 1.276 ($\pm 0.404$) | 0.354 ($\pm 0.116$) | 0.360 ($\pm 0.127$) |
| **SVR-RBF** | Descriptors | GEx | 237 | 0.680 ($\pm 0.110$) | 1.278 ($\pm 0.403$) | 0.348 ($\pm 0.120$) | 0.354 ($\pm 0.135$) |
| **SVR-Linear** | Descriptors | GEx | 237 | 0.666 ($\pm 0.102$) | 1.292 ($\pm 0.420$) | 0.324 ($\pm 0.119$) | 0.331 ($\pm 0.126$) |
| **SVR-Linear (w/RFE)** | Descriptors | GEx | 237 | 0.666 ($\pm 0.102$) | 1.293 ($\pm 0.421$) | 0.322 ($\pm 0.118$) | 0.330 ($\pm 0.124$) |
| **PathDNN** | Drug Targets | GEx, pathway information | 234* | 0.612 ($\pm 0.074$) | 2.201 ($\pm 0.698$) | 0.193 ($\pm 0.061$) | 0.170 ($\pm 0.078$) |
| **tCNN** | SMILES one-hot encoding | Genetic Features | 237 | 0.586 ($\pm 0.060$) | 1.369 ($\pm 0.427$) | 0.147 ($\pm 0.068$) | 0.147 ($\pm 0.072$) |

Based on these evaluations, BiG-DRP+ has the best performance using all metrics, while BiG-DRP has the second-best performance. The BiG-DRP+ clearly outperforms MLP, further highlighting the importance of the bipartite graph and H-GCN in the drug response prediction task. Similar to LPO-CV evaluation, a drug-wise analysis using SCC for each drug showed a significantly superior performance of BiG-DRP+ compared to all baseline methods (one-sided Wilcoxon Signed-Rank test, Supplementary Table S2). Supplementary Table S1 provides the drug-specific performance metrics for all drugs.

To assess the generalizability of BiG-DRP+ to independent datasets, we used it to predict the drug response of patient tumours from the TCGA dataset treated with cisplatin, gemcitabine, doxorubicin, and paclitaxel. Given the predicted log IC50 values, we used a one-sided statistical test to determine if our models can distinguish between the patients that are resistant from those that are sensitive to these two drugs (using all TCGA samples with known clinical drug response). Our statistical analysis (Mann Whitney U test, since data corresponding to one of the drugs did not pass test of normality) showed significant p-values for three drugs (p = 2.19E-7 for cisplatin, p = 8.80E-3 for doxorubicin, and p = 3.40E-2 for gemcitabine). Next, we removed any tumour sample that had received a different drug beforehand or during the period that our drug of interest was administered. Even though this significantly reduced the number of samples, the results (Welch’s t-test, since data corresponding to all drugs passed test of normality) were significant for cisplatin (p = 1.82E-2) and doxorubicin (p = 4.29E-2). Supplementary Table S3 provides detailed information regarding the samples and the results of different statistical tests.

### Detailed Evaluation of BiG-DRP+

Since one major component of the BiG-DRP and BiG-DRP+ pipeline is the bipartite graph of the CCLs and drugs, we sought to evaluate the effect of different thresholds for forming this graph. As explained in Methods, a drug is connected to a CCL with a sensitive (resistant) edge if the log IC50 of the CCL is among the bottom (top) k% of all the CCLs. In our analysis, we fixed this value to be k = 1. To assess the robustness of the results to this parameter, we formed different bipartite graphs with different choices of k = 0.5, 1, 2, 5, 10 and repeated the LPO-CV and LCO-CV. Supplementary Table S4 provides the SCC and AUROC of BiG-DRP and BiG-DRP+ for these evaluations for different values of k. These results suggest that the performance of our proposed methods remain stable for these different choices of k, with a slight deterioration as k increases (less than 1% in all evaluations when comparing k=10 to k=1). This deterioration is expected, since an increase in k increases potentially erroneous edges in the bipartite graph.

Next, we asked whether the choice of drug features as attributes in the bipartite graph has a significant effect on the performance of BiG-DRP+. To address this question, we used Morgan fingerprints [13] of the drugs, alone or in addition to the drug descriptors, as the attributes of the drug nodes in the bipartite graph. The results (Table 3) revealed that there is not a substantial difference between any of these choices, but simultaneously using both types of drug features slightly improves the results.

**Table 3:** The performance of BiG-DRP+ with different drug attributes. The rows show the results of BiG-DRP+ when drug descriptors (vectors of length 198), Morgan fingerprints (vectors of length 512), or the combination of both (vectors of length 710) are used as node attributes.

| Method | Drug Attribute | LPO-CV |  | LCO-CV |  |
| --- | --- | --- | --- | --- | --- |
| | | AUROC mean (± std.) | SCC mean (± std.) | AUROC mean (± std.) | SCC mean (± std.) |
| BiG-DRP+ | Descriptors | 0.878 (±0.068) | 0.748 (±0.100) | 0.746 (±0.077) | 0.431 (±0.094) |
| | Morgan | 0.878 (±0.068) | 0.748 (±0.100) | 0.743 (±0.080) | 0.426 (±0.098) |
| | Both | 0.879 (±0.068) | 0.748 (±0.099) | 0.746 (±0.077) | 0.433 (±0.095) |

Finally, we asked how different choices of hyperparameters influence the performance of BiG-DRP+. For this purpose, we ran our model with 648 different combinations of learning rate (5E-5, **1E-4**, 5E-4, 1E-3), batch size (64, **128**, 256), CCL encoder size (512, **1024**, 2048), H-GCN size (256, **512**, 1024), predictor hidden layer size (256, **512**, 1024), and dropout (**with** or without). (The bold-face options represent the default values used for our models). The stacked histogram in Figure 3A shows the mean SCC value of these combinations in a 5-fold LCO-CV framework. Interestingly, there are 82 combinations that perform on par with the default parameters and 47 combinations that perform better. This suggests that if computational complexity is not of a concern, one may improve the performance of BiG-DRP+ by tuning the hyperparameters.

**Figure 3:**
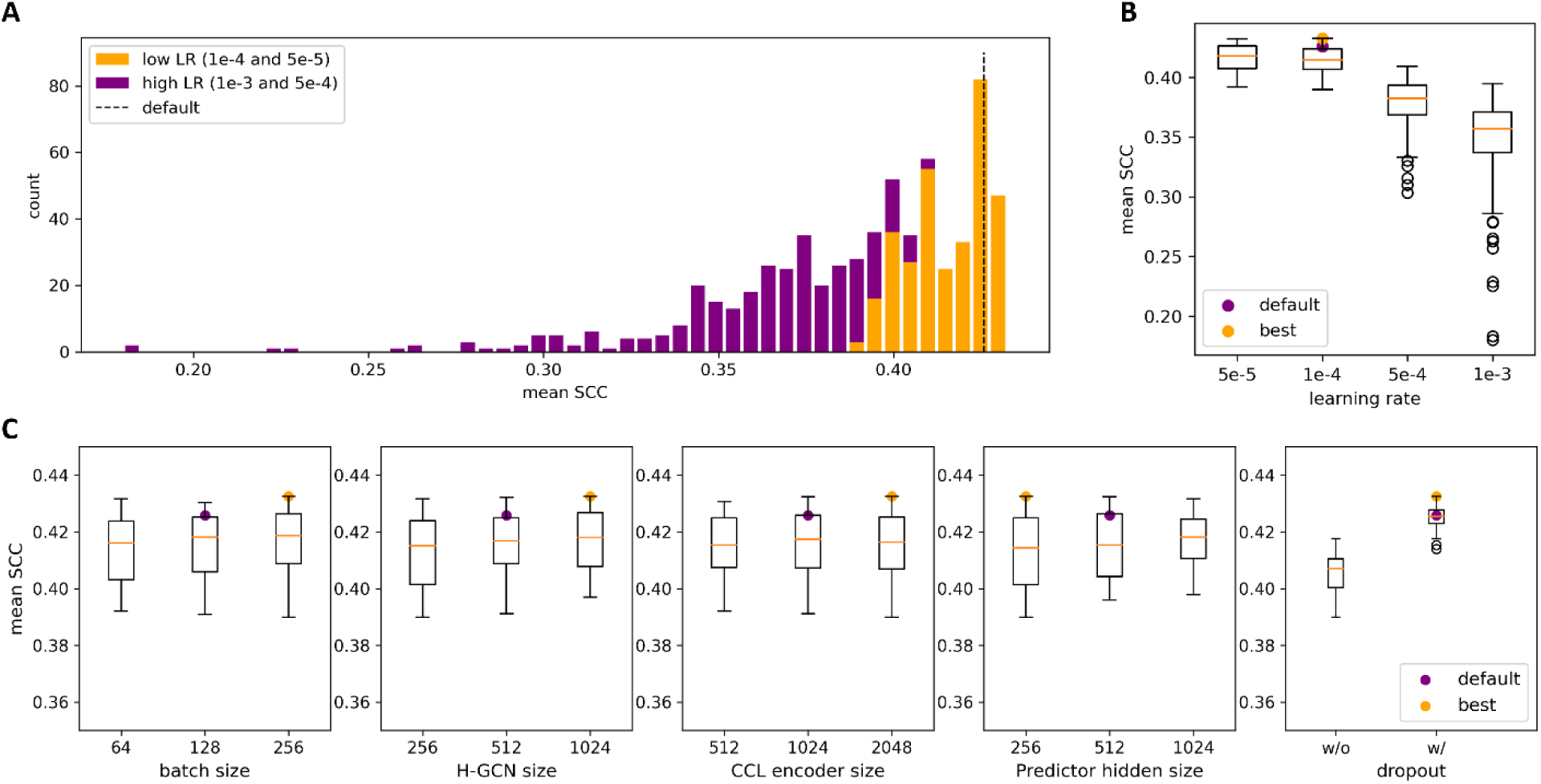
Performance comparison of BiG-DRP for different combinations of hyperparameters. A) The distribution of mean SCCs of the models in 5-fold LCO-CV. The colors correspond to the fraction of the bin that utilized either a high or low learning rate. B) The boxplots of the mean SCCs grouped by the learning rate. C) The boxplots of the mean SCCs for combinations that used 1E-4 and 5E-5 as learning rates, grouped by specific hyperparameters. The boxes range from the first to the third quartile, while the horizonal line corresponds to the median. The purple datapoint represents the default hyperparameter combinations and the orange datapoint pertains to the combination of hyperparameters that performed best.

Further analysis revealed that the learning rate is the most influential hyperparameter (Figure 3B) and a relatively large learning rate deteriorates the performance; however, learning rates of 5E-5 or 1E-4 (the default) work well. More importantly, if the learning rate is selected appropriately (the two choices mentioned above), the effect of other hyperparameters is relatively small and the majority of choices result in good performance (orange fraction of the histogram in Figure 3A). Figure 3C better illustrates this by depicting the mean SCC for different choices of hyperparameters when only learning rates of 5E-5 and 1E-4 are included. The only other hyperparameter that seems to play an important role is dropout, where its inclusion (slightly) improves the performance.

### Characterization of the bipartite graph

Next, we sought to better characterize the bipartite graph and the drugs that have most benefited from using this graph in the drug response prediction task. For this purpose, we first formed a single bipartite graph by aggregating the bipartite graphs corresponding to each of the five folds in our LCO-CV evaluation (i.e., by finding the union of edges). Then, we used a nested stochastic block model (NSBM) [39] to infer the modular substructure of the graph, while taking into account the edge type (i.e., resistant and sensitive) connecting each two nodes. This approach automatically identifies the number of clusters by maximizing the likelihood of the graph being generated from the partitioning. The final partitioning is based on running the stochastic algorithm many times (in our case 1000 times) and selecting the number of clusters and the partitioning that is most frequently supported by these runs. The number of clusters varied between 17 to 20 (Figure 4A), with 18 selected by the algorithm as the final number of clusters (5 drug clusters and 13 CCL clusters). Comparing the clusters identified by each run of the algorithm with the final clusters using Rand Index (RI) [40] revealed a high degree of concordance (Figure 4B, mean RI = 0.89 ± 0.01).

**Figure 4:**
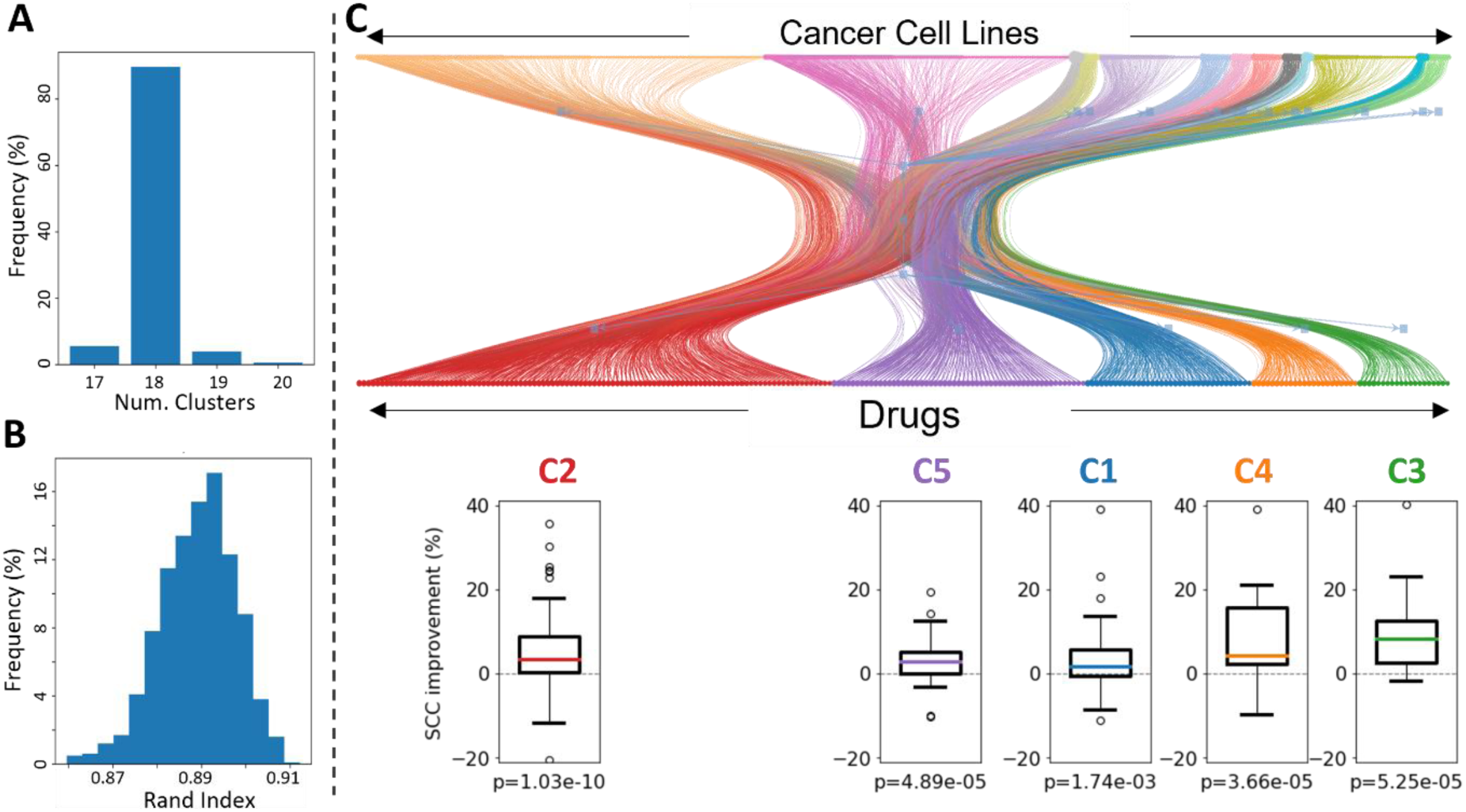
The CCL-drug bipartite graph and its clusters. A) The histogram shows the number of clusters obtained by NSBM in each run (total of 1000 runs). B) The histogram shows the Rand Index between each clustering (in each run) with the final cluster assignment. C) The graph represents the bipartite graph and the boxplots show the distribution of SCC improvements obtained for each drug using BiG-DRP+ compared to MLP in the LCO evaluation. The p-values are obtained using a one-sided Wilcoxon signed rank test.

Figure 4C illustrates the bipartite graph and clusters identified using this method (see Supplementary Table S5 for the cluster assignment of drugs and CCLs). In particular, five drug clusters were identified. Comparing the performance of BiG-DRP+ compared to MLP (SCC-LCO), revealed that all these clusters significantly benefit from the use of the bipartite graph (one-sided Wilcoxon signed rank test, Figure 4B). In particular, Cluster 3 had the highest median improvement in SCC (8.4%) and had a significant improvement p-value (p = 5.25 E-5). The majority of the drugs in this cluster (13 out of 20) are protein kinase inhibitors, with 8 of them targeting members of serine/threonine protein kinase family and 5 of them targeting members of tyrosine kinase family. These observations suggest that information sharing across the bipartite graph used in our methods benefit certain groups of drugs more than others and this may be dependent on the similarity between drugs’ mechanisms of action.

Next, we sought to characterize the CCL clusters. Supplementary Table S6 shows the enrichment of CCL clusters in tissue type, cancer type, and driver mutations (hypergeometric test, corrected for multiple tests using Benjamini-Hochberg FDR). The analysis revealed that while only two clusters (out of 13) were enriched in cancer type (FDR < 0.05), namely cluster 1 in B-Lymphoblastic Leukemia and cluster 4 in Chronic Myelogenous Leukemia, the majority of clusters (9 out of 13) were enriched in at least one driver gene mutation. For example, cluster 1 was enriched in CCLs with mutations in RBM38 and GNA13, while cluster 2 was enriched in CCLs with mutations in POLQ and BRCA1. These observations suggest that the patterns captured by the bipartite graph goes beyond tissue or cancer types and is able to capture patterns at the molecular level.

### Identification of biomarkers of drug sensitivity

To identify genes whose expression substantially contribute to the predictive model, we used a pipeline similar to the one we proposed in a previous study [5]. This approach provides an aggregate contribution score for each gene in the model and uses these scores to systematically identify the set of top contributing genes in each model. We focused on 15 drugs for which BiG-DRP+ provided the highest SCC values in the LCO-CV evaluation. Supplementary Table S7 provides the ranked list of genes that were implicated for each of the 15 drugs. We clustered the drugs based on the contribution scores of all implicated genes (Figure 5). Interestingly, four drugs formed a clear cluster, separate from the others: trametinib, refametinib, selumetinib, and pd0325901. Further investigation revealed that these drugs all are MEK inhibitors (i.e., inhibit the mitogen-activated protein kinase kinase enzymes) and involve some similar mechanisms of action [10].

**Figure 5:**
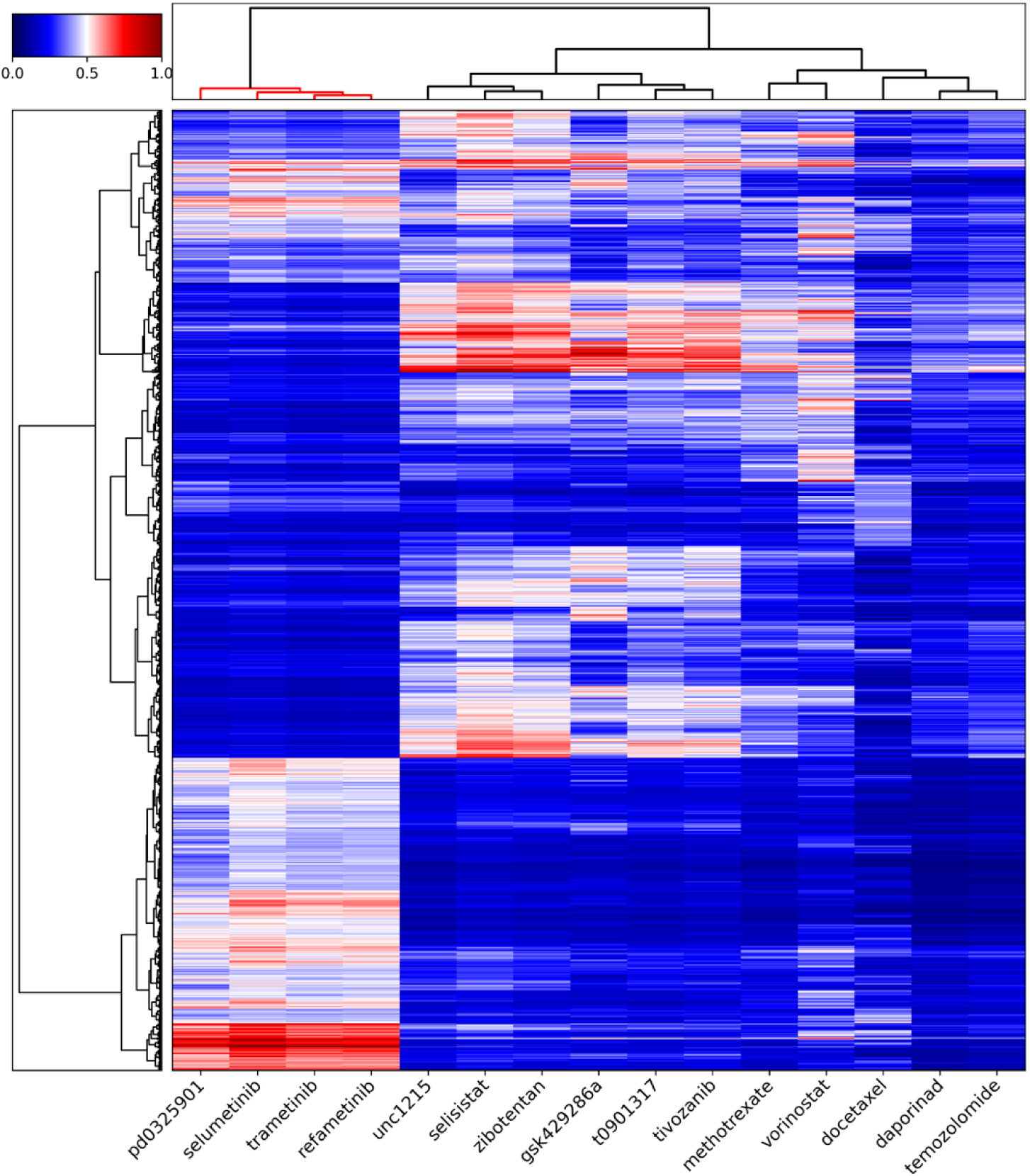
The clustering of 15 drugs based on contribution scores of their genes. The contribution scores of the union of genes implicated for these drugs is used to cluster drugs using hierarchical clustering. The heatmap shows the contribution scores.

Next, we focused on genes implicated for trametinib, a MEK-inhibitor for which BiG-DRP+ had the best performance (SCC in LCO-CV). For this drug, *ETV5* had the highest prediction contribution. ETV5 and ETV4 (the fourth highest contributor) are among the ETS family of oncogenic transcription factors. The expression of this family has been shown to be upregulated in solid tumours and they have been shown to be involved in tumour’s progression, metastasis and chemoresistance [41]. Previous studies have shown ETV5 to be regulated by ALK, a receptor tyrosine kinase, in a MEK/ERK-dependent manner in neuroblastoma cell lines [42]. In addition, treatment of various cancer cell lines with trametinib has been shown to downregulate ETV5 [42–44]. Moreover, the overexpression of ETV4 and ETV5 have been shown to reduce the sensitivity different cancer cell lines to this drug [44].

To obtain a better functional characteristic of the genes implicated for trametinib, we also performed pathway enrichment analysis on genes implicated for this drug (see Supplementary Table S8 for results of pathway enrichment analysis of all 15 drugs). Several important pathways related to MAPK signaling, EGFR signaling, and IL2 signaling were identified (Fisher’s exact test, FDR<0.05). Taken together, these results suggest that genes that contribute to the predictive ability of BiG-DRP+ for trametinib point to important genes and signaling pathways involved in its mechanism of action.

### Mutation landscape of TCGA tumor samples and their association with drug response

Next, we sought to evaluate the mutation landscape of tumors in TCGA dataset and their associations with drug response predicted using Big-DRP+. For this purpose, we predicted the normalized logIC50 of 9067 TCGA tumors (that had both mutation and GEx data) corresponding to 32 cancer types to 237 drugs in our training dataset (Supplementary Table S9, Methods). We identified 10 genes that were mutated in more than 10% of the samples (Supplementary Table S9) and performed two-sided Mann–Whitney U tests to assess the association between mutations in these genes and drug response (the FDR values reported in this study correspond to this test). In this section we focus on the insights obtained from *PIK3CA* mutation due to its important role in determining the drug response in various cancers and its potential as a therapeutic target [45] (results of statistical tests for all genes are provided in Supplementary Table S9).

*PIK3CA* is an oncogene whose mutation leads to hyperactivation of PI3K/AKT/mTOR pathway, associated with cancer progression and poor outcome in many cancer types [46–49]. Various targeted therapies have been developed to target and inhibit this pathway in patients with deregulation and hyperactivity of PI3K/AKT/mTOR pathway (due to *PIK3CA* mutation or other mechanisms such as loss or inactivation of *PTEN*) [50]. Additionally, various studies have shown that mutation in this gene is associated with better response to PI3K inhibitors both *in vitro* and *in vivo* [50, 51]. Consistent with these, our pan-cancer analyses showed that tumors that harbor this mutation are significantly more sensitive to drugs targeting PI3K/AKT/mTOR pathway (Figure 6A, one-sided Wilcoxon signed rank test P = 1.14E-5) such as the pan-AKT kinase inhibitor GSK690693 (FDR = 2.13E-59) and the pan-class I PI3K inhibitor ZSTK474 (FDR = 1.15E-29) (Supplementary Table S9).

**Figure 6:**
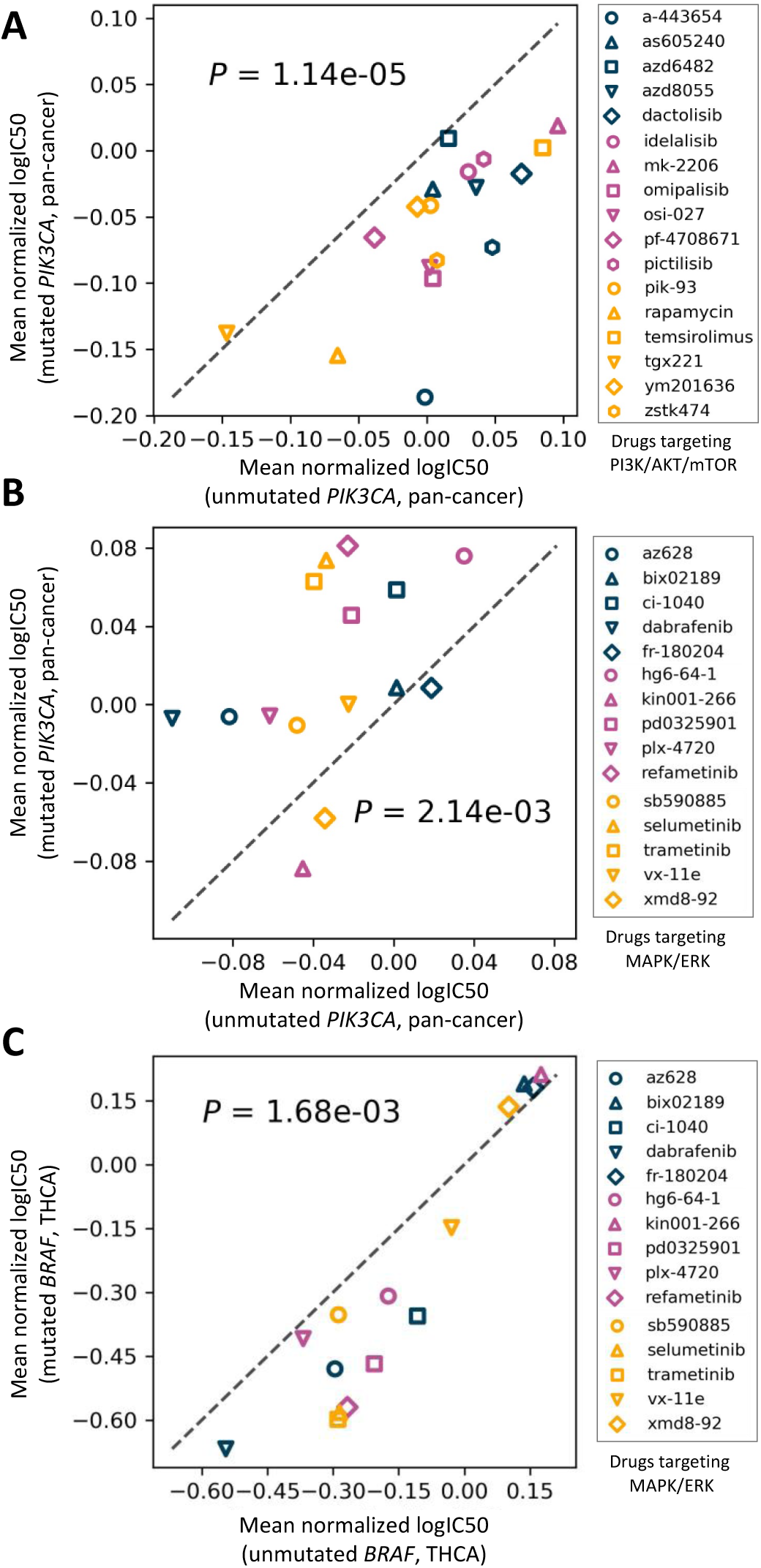
The association between mutations and drug response in TCGA. The scatter plots show the mean predicted normalized logIC50 for mutated and unmutated tumors. P-values are calculated using a one-sided Wilcoxon signed rank test. A) The association between *PIK3CA* mutation and response to drugs targeting the PI3K/AKT/mTOR pathway in our pan-cancer study. B) The association between *PIK3CA* mutation and response to drugs targeting the MAPK/ERK pathway in our pan-cancer study. C) The association between *BRAF* mutation and response to drugs targeting the MAPK/ERK pathway in THCA.

On the other hand, mutation in this gene was associated with increase in resistance to drugs targeting the MAPK/ERK signaling pathway (Supplementary Table S9). In particular, the mean predicted normalized logIC50 of *PIK3CA*-mutated tumors were significantly larger for drugs targeting this pathway compared to tumors that did not harbor this mutation (one-sided Wilcoxon signed ranked test P = 2.14E-3, Figure 6B). Various studies (both *in vivo* and *in vitro)* have shown a regulatory link between MAPK/ERK and PI3K/AKT/mTOR pathways and inhibition of MAPK/ERK signaling has been linked to an increase in the activity of PI3K/AKT/mTOR pathway ([52] and references therein). Previous studies have shown that hyperactivity in PI3K/AKT/mTOR pathway as a result of *PIK3CA* mutation increases drug resistance to dabrafenib and trametinib (drugs targeting MAPK/ERK pathway), supporting our observations (dabrafenib FDR = 2.93E-18, trametinib FDR = 7.64E-9, Supplementary Table S9). Mutation in *PIK3CA* has been shown to confer resistance to PD0325901 [53], a MEK-inhibitor that decreases MAPK/ERK pathway activity, and genetic ablation of the mutant allele of this gene has been shown to increase sensitivity to this drug in MEK-inhibitor resistant cells [53]. Our analysis also showed that tumors harboring *PIK3CA* mutation are more resistant to this drug (FDR = 1.33E-5).

Among the four drugs that target IGF1R, three showed a significantly higher predicted logIC50 value in *PIK3CA*-mutated tumors. Previous studies have shown a link between this protein and *PIK3CA-*driven ovarian cancer [54] and breast cancer tumors harboring mutation in this gene [55], suggesting the dual inhibition of PI3K and IGF1R as a new therapeutic approach. Another noteworthy example identified by our analyses is cetuximab (FDR = 5.98E-3), which is an epidermal growth factor receptor inhibitor. Previous studies have shown an association between activity of PI3K/AKT/mTOR pathway and resistance to this drug [56].

To assess the effect of mutations on drug resistance in a cancer type-specific manner, we focused on thyroid carcinoma (THCA), which is the most common endocrine malignancy, as an illustrating example [57]. In this cancer type, only *BRAF* was mutated in more than 10% of the samples (mutated in 57.7% of tumors). The mutation in this gene, the most frequent of which in thyroid cancer is V600E mutation [57, 58], activates the MAPK/ERK pathway resulting in sustained cell proliferation adverse phenotypes [57]. Various studies have proposed this pathway as a therapeutic target, and have shown that cancer cells (including those corresponding to thyroid cancers) harboring this mutation are much more sensitive to BRAF-inhibitors (e.g., AZ628 [59]) and various MEK-inhibitors [60]. Our analyses also showed that THCA tumors harboring *BRAF* mutation are significantly more sensitive to drugs targeting MAPK/ERK pathway (Figure 6C, one-sided Wilcoxon signed rank test P = 1.68E-3), including BRAF-inhibitors AZ628 (FDR = 2.25E-21) and HG6-64-1 (FDR = 5.62E-12), and MEK-inhibitors such as trametinib (FDR = 2.83E-26), refametinib (FDR = 1.69E-25), and selumetinib (FDR = 1.75E-25).

Taken together, these results suggest the utility of our proposed model in providing insights in pharmacogenomics studies.

## DISCUSSION AND CONCLUSION

In this study, we proposed two novel graph representation deep learning methods to incorporate information regarding the sensitivity and resistance of cell lines, their gene expression profiles and chemical drug attributes to obtain better drug representations. Using cross-validation and different data splitting methods we showed significant improvement compared to traditional and state-of-the-art methods. Using a computational pipeline to make neural networks explainable, we identified a set of genes that substantially contribute to the predictive model. These genes implicated important signaling pathways and pointed to shared and unique mechanisms of action in the drugs. In addition, we performed a study on the association between the mutation status of cancer tumors from TCGA and the predicted drug response. These analyses revealed various insights, many of which were confirmed by independent studies, which further illustrates the utility of our pipeline in pharmacogenomics studies.

Moreover, detailed evaluation of our methods showed a high degree of robustness towards changes in the threshold used to form the bipartite graph. This further supports the importance of different techniques we used to ensure stability of our proposed architecture: the normalization factor and the injected self-loop in our H-GCN’s forward pass. More specifically, due to the injected self-loop, the nodes retain a portion of their own information, which forces the embeddings to have some level of separation. The normalization factor also helps by preventing the received messages from becoming too large and overpowering the self-loop. It is important to note that this robustness may not be applicable to some corner cases. For example, when a drug’s connected CCLs are not connected to any other drug (i.e., it forms a disconnected star subgraph), this drug’s embedding would not benefit from the existence of the second H-GCN layer. As another example, the second H-GCN layer will be obsolete if all the drugs happen to form disconnected stars, and thus no information sharing will take place across drugs. Another example is when we add a new drug that results in a disconnected node. A disconnected node will not be able to incorporate CCL information into the drug embedding, which defeats the purpose of the H-GCN.

Unlike many previous models (e.g., NRL2DRP [30]) that require both cell lines and drugs to be present in the training set, BiG-DRP is designed to enable prediction of unseen cell lines (those that are not present in the training set). However, the drug embedding part of the model (the H-GCN) requires the drugs to be part of the bipartite graph. This constraint implies that the drugs present in the test set must be also present in the training set. As a result, this model generally is not applicable to predict the response of CCLs to unseen new drugs. Although this could be naively remedied by assuming known edges involving the unseen drug in the bipartite graph, this kind of solution is impractical and would be difficult to enact without reducing the test set. However, in most practical applications (e.g., prediction of drug response of cancer patients [4] and [5]), it is more crucial for the model to generalize to unseen samples (CCLs or patients). The reason is that before a drug enters clinical trial or enters clinical usage, many *in vitro* studies on CCLs are first performed. Consequently, one can expect to have access to molecular description and drug response of a drug for which the drug responses of a new set of samples (CCLs or patients) are to be predicted.

In this study, instead of directly using the logIC50 of drugs, we normalized the logIC50 of each drug (separately) across the CCLs. This was done first to ensure that our prediction performance results are not artificially inflated and second to make the drug response ranges of different drugs comparable to allow the model to learn useful representations across drugs. However, this normalization means that the predicted values should not be used to compare the potency of different drugs, but rather should be used to compare the sensitivity of different CCLs to a specific drug. This is why when we reported our prediction performance results, we calculated them one drug at a time (across CCLs). If one wants to recover logIC50 values, these predictions can be easily modified to reverse the normalization and allow comparison of different drugs for the same CCL.

One of the main motivations of this study was to improve the representations of drugs for the task of drug response prediction. While direct drug targets or SMILES chemical information of drugs are common approaches for representing drugs, we believe these representations can be improved by capturing the effects these drugs have on CCLs, either by measuring the changes in the GEx profiles of CCLs after administration of the drug (e.g., LINCS dataset [61]) or using the bipartite graph formulation proposed in this study. Improved drug representations are particularly important in more challenging tasks such as prediction of response to drug combinations, in which the sheer number of possible drug combinations (even for drug-pairs) means that experimental measurements can only capture a very small portion of all possibilities. As a result of this small sample size problem, more informative and robust drug representations become crucial in developing generalizable machine learning models for drug combinations, a direction that we will pursue in the future by generalizing the models introduced in this study.

## Authors’ contributions

AE and DEH conceived the study, designed the project and the algorithms. DEH implemented the pipeline. DEH and YL ran the baseline methods and performed the statistical analyses of the results. All authors read and approved the final manuscript.

## SUPPLEMENTARY TABLES

**Supplementary Table S1**: The performance of BiG-DRP+, BiG-DRP and baseline methods (columns) using five-fold CV for each drug (rows). Values presented are the mean across five folds. Each tab corresponds to a performance metric (Spearman’s correlation coefficient and area under the receiver operating characteristic) for each data-splitting scenario: leave-cell lines-out (LCO) and leave-pairs-out (LPO).

**Supplementary Table S2**: The results of the one-sided Wilcoxon signed rank test, represented by the p-values. The test compares the per-drug Spearman’s correlation coefficient (SCC) of the BiG-DRP+ and the other methods, where the alternative is that BiG-DRP+’s SCC is significantly larger.

**Supplementary Table S3:** The description of samples and results of statistical tests for prediction of clinical drug response of TCGA cancer patients.

**Supplementary Table S4**: The performance of BiG-DRP and BiG-DRP+ using for *k* ∈ {0.5, 1, 2, 5, 10}. The performance metrics were calculated independently per drug and presented as the mean and standard deviation.

**Supplementary Table S5**: The cluster assignments of the aggregated bipartite graph using the nested stochastic block model. The first tab shows that cluster indices for the drugs and the second tab shows the cluster indices for the cell lines. Although the graph partitioning was performed on the entire graph, drug and cell line clusters were mutually exclusive; as a result, similar indices in different tabs do not pertain to the same cluster.

**Supplementary Table S6:** Characterization of CCL clusters in the bipartite graph with respect to tissue type, cancer type and driver mutations.

**Supplementary Table S7**: The top genes and their normalized contribution scores for top-performing drugs in the leave-cell lines-out scenario. Each tab corresponds to the top genes for a specific drug.

**Supplementary Table S8**: The results of the pathway enrichment analysis of the top genes (Supplementary Table S7) on the Reactome pathways using Fisher’s exact test. Each tab corresponds to a specific drug. The corrected p-values are indicated in the *pvalue_cor* column.

**Supplementary Table S9:** The association between predicted drug response and mutation status of TCGA samples for our pan-cancer and THCA study.

